# A Knock-In *Igfn1^iCre^* transgenic mouse line provides partial developmental access to type-7 bipolar cells

**DOI:** 10.64898/2026.03.06.710004

**Authors:** Shambhavi Chaturvedi, Haruka Yamamoto, Akihiro Matsumoto, Manabu Abe, Toshikuni Sasaoka, Keisuke Yonehara

**Affiliations:** Multiscale Sensory Structure Laboratory, National Institute of Genetics, 1111 Yata, Mishima, 411-8540, Shizuoka, Japan; The Graduate Institute for Advanced Studies, SOKENDAI, Shonan- Kokusai-Mura, Hayama, 240-0115, Kanagawa, Japan; Department of Animal Model Development, Brain Research Institute, Niigata University, 1 Bancho-757 Asahimachidori, Chuo Ward, Niigata, 951-8122, Japan; Department of Comparative and Experimental Medicine, Brain Research Institute, Niigata University, 1 Bancho-757 Asahimachidori, Chuo Ward, Niigata, 951-8122, Japan; Danish Research Institute of Translational Neuroscience–Nordic-EMBL Partnership for Molecular Medicine, Department of Biomedicine, Aarhus University, Aarhus C, DK8000, Denmark

**Keywords:** Bipolar cell subtypes, Development, Retina, Genetic labelling

## Abstract

Functional neuronal circuits in the vertebrate retina emerge through coordinated developmental events, yet the timeline by which bipolar cells acquire visual feature selectivity remains unclear. A major barrier is the limited genetic access to bipolar subtypes during early postnatal stages. Recent comprehensive transcriptomic study points to *Igfn1* as a molecular marker for type-7 bipolar cells (BC 7), a subtype that exhibits direction-selective glutamate releases in adult. Here, we generated an *Igfn1*^*iCre*^ knock-in mouse line and characterized *Igfn1*–positive cell morphology from postnatal day 4(P4) to adult using Cre-dependent tdTomato reporter mice. We found *Igfn1*–positive cells in the inner retina by P12–P15, predominantly labelling bipolar cells and some amacrine populations. At P15, about 71% of labelled bipolar cells stratified their axons in the S4 sublamina of the inner plexiform layer, consistent with BC 7 morphology. In adult retina, the widespread *Igfn1*–labelling appears slightly dominated in amacrine cells. To validate these observations, we analysed *Igfn1* expression in the Mouse Retina Cell Atlas and confirmed strong *Igfn1* enrichment in BC 7 and expression in additional retinal cell types, mirroring experimental results. Overall, these results reveal *Igfn1*^*iCre*^ as a potential developmental tool for BC 7 access in the retina.

## 2. Introduction

The vertebrate inner retina comprises a layered network of excitatory and inhibitory interneurons that transform outer photoreceptor signals into parallel channels to highly specialized retinal ganglion cells (Hsiang et al. (2024)). Among these interneurons, bipolar cells provide the principal excitatory relay by integrating signals from light sensitive photoreceptors to the output neurons (Euler et al. (2014)). Mammalian bipolar cells are subdivided into more than a dozen molecularly, morphologically and physiologically distinct types, each connected to defined amacrine and ganglion cells to support complex visual computations (Franke et al. (2017); Shekhar et al. (2016)). One such computation is motion direction selectivity, the ability to preferentially respond to stimuli moving in specific directions (Matsumoto et al. (2021)). Recent work has identified type-2 and type-7 bipolar cells as key contributors in this computation (Matsumoto et al. (2021)). In this study, we focus on type-7 bipolar cells (BC 7). In the adult retina, BC 7 are best identified by their distinctive axon morphology, with terminals narrowly stratifying in sublamina S4 of the Inner Plexiform Layer (IPL), proximal to the inner starburst amacrine cell (SAC) band (Wässle et al. (2009); Stabio et al. (2018); Hall et al. (2019)). Functionally, BC 7 are ON bipolar cells, depolarizing in response to bright stimuli positioned in the receptive field center.

The developmental onset and molecular mechanisms underlying direction selectivity in BC 7 remain elusive. A major limitation has been the lack of subtype-specific genetic access during postnatal development stages, when functional properties are established. Available genetic mouse lines co-label multiple bipolar subtypes and therefore lack the specificity required to selectively target BC 7. For example, Grm6-Cre drives recombination across all ON bipolar cells (Ueno et al. (2018)), and Gus-GFP labels both rod and ON cone bipolar cells (Huang et al. (2003)).

To address this limitation, we referred to latest comprehensive single-cell transcriptomic analyses in the adult retina as a path forward. Shekhar et al. (2016) report *Igfn1* (immunoglobulin-like and fibronectin type III domain containing 1) as a BC 7-enriched marker. Additional transcriptomic studies suggest no representation of *Igfn1* as a marker to any amacrine or ganglion cell classes (Yan et al. (2020); Goetz et al. (2022)). Interestingly, an important support for its developmental relevance comes from birthdating experiments that used a SABER-FISH strategy to identify bipolar subtype genesis (West and Cepko (2022)). The study demonstrates *Igfn1* expression at P18 in the inner retina, where cell bodies of bipolar cells and amacrine cells are located. On this basis, we postulated that an *Igfn1*-driven recombinase could provide genetic access to BC 7. Yet no *Igfn1*-based driver line had been reported, and whether *Igfn1* maintains bipolar cell specific subtype restriction across development was unknown.

Here, to genetically access the BC 7, we generated a knock-in *Igfn1*^*iCre*^ mouse line and characterized the Cre-expression using a tdTomato reporter mice. Our results show selective access to BC 7 compared to other bipolar cells around P15, a developmental stage for which subtype-specific tools were previously not available. However, the genetic access is not completely exclusive as smaller fraction of additional bipolar and amacrine populations are also labelled. In adult stage, we observed widespread labelling in the inner retina which was slightly more dominant in amacrine cells. To validate these observations, we examined *Igfn1* expression in the publicly available Mouse Retina Cell Atlas (MRCA) (Li et al. (2024)). This analysis confirmed BC 7 enrichment with comparatively lower expression in other retinal types, mirroring experimental findings. To expand the utility of the transgenic mouse line, we characterized the mouse brain at the adult stage. We found prominent access to anterior forebrain brain regions, including cortical and hippocampus neurons. Overall, these results demonstrate the *Igfn1*-based driver line as a tool that provides partial genetic access to BC 7 and other inner retinal and brain neurons while opening the question how to selectively target the bipolar subtype in adult.

## 3. Methods

### a. Experimental model and subject details

*Igfn1*^*iCre*^ mouse line was generated at Niigata university in a C57BL/6N background. The mice were generated using BAC clone RP23-192B8 and RP23-320P13 encoding the mouse *Igfn1*. The coding sequence at first ATG sequence of exon1 was replaced by iCre-splice gene cassette and neomycin resistant gene. The insertion cassette encodes a codon-improved Cre recombinase containing an inserted splice intron derived from the SV40 t-antigen gene, positioned between codons 283 and 284, as previously described (Inoue et al. (2018)). The BAC recombinants were electroporated first into ES cells and upon positive selection by drug were implanted into adult female C57BL/6N mice. The transgene insertion was determined by PCR using iCre recombinase primers, by amplification of 371bp fragments. Finally, the knock-in mice were backcrossed to C57BL/6J mice. All mice were maintained in the animal facility of the National Institute of Genetics. Mice were kept in conventional cages under standard controlled conditions (temperature 23 ± 2°C, humidity 50 ± 10%, 12 h light-dark cycle) with ad libitum access to food and water. All animal experiments were performed according to standard ethical guidelines and were approved by the Institutional Animal Committees (Reference Number: No. R7-1).

The heterozygous or homozygous Cre-mouse line were crossed with homozygous Rosa-STOP-tdTomato reporter mice for cell morphology imaging. The day on which pups were born were considered as postnatal day 0 (P0). Further, P4 to adult (P28 or above) week-old mice of either sex were used for experiments.

### b. Histochemistry

#### Retina samples

The mice were sacrificed by cervical dislocation or isoflurane. From each mouse, two eye ball were isolated and both retinae were dissected. Next, retinae were fixed for 30 min in 4% paraformaldehyde (PFA) in phosphate buffer saline (PBS) and washed with PBS at room temperature (RT). For whole mount staining, the retinae were first incubated in 30% sucrose in PBS overnight at 4°C. To enhance the penetration of antibodies, retinae were frozen and thawed three times in the same sucrose buffer. After washing with PBS, retinae were blocked for 1-2 h in blocking buffer (1% bovine serum albumin [BSA], 10% normal donkey serum [NDS], 0.5% Triton X-100, 0.02% sodium azide in PBS) at RT. The retinae were incubated with primary antibodies as mentioned in Table 1 for 5 days at RT in antibody reaction buffer (1% BSA, 3% NDS, 0.5% Triton X-100, 0.02% sodium azide in PBS), and secondary antibodies (refer Table 1) overnight at 4°C in antibody reaction buffer. The fluorescent samples were counterstained with DAPI (4’,6-diamidino-2-phenylindole, Fujifilm Wako). After a final washing in PBS, retinae were mounted on glass slides (Matsunami Glass) and embedded with ProLong™ Glass Antifade Mountant (ThermoFisher Scientific).

**Table 1.**
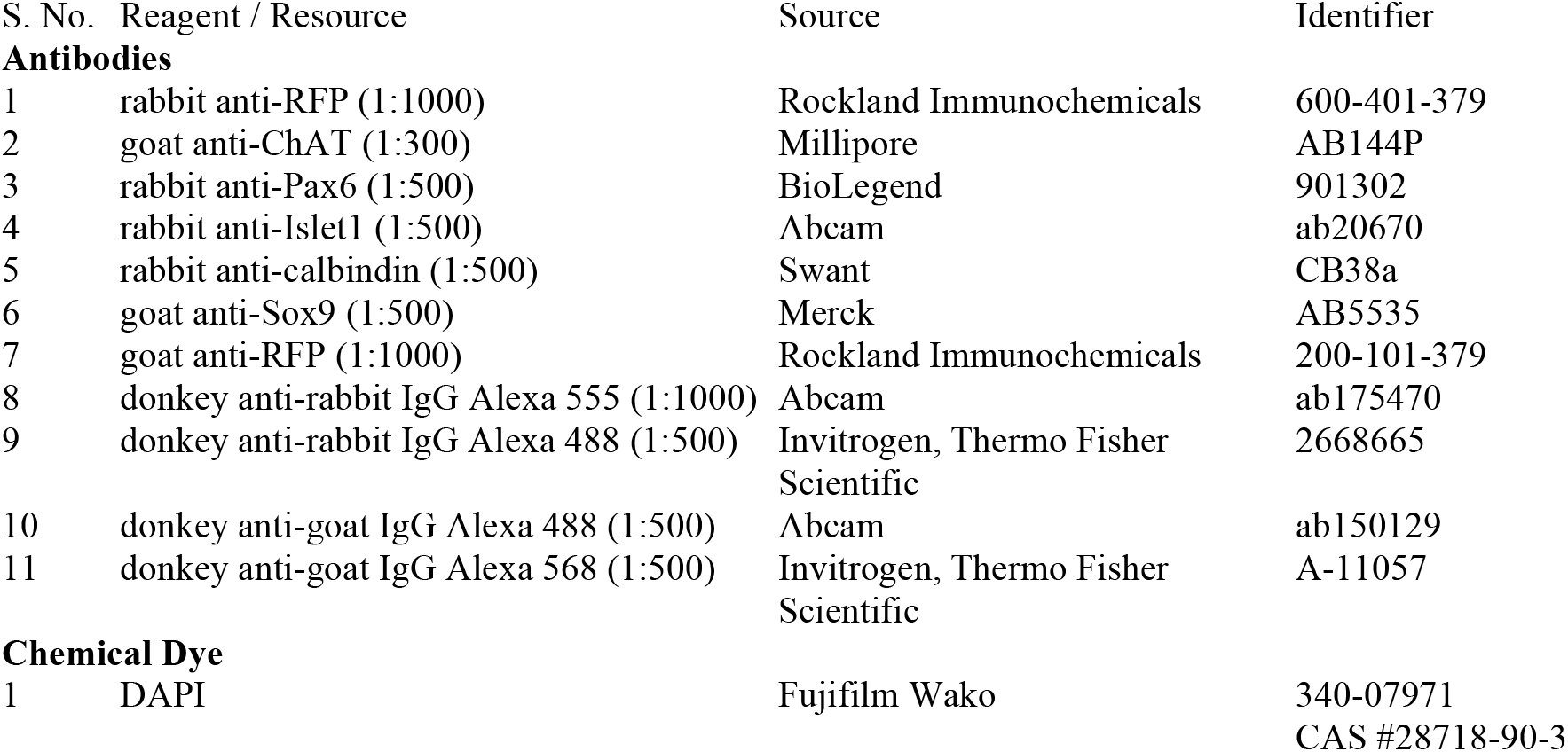
Reagents and resources used in this study.

To prepare cryosections, retinae were immersed in 30% sucrose in PBS for 2 days at 4°C and frozen in OCT compound (Sakura Finetek). Vertical sections of 20-*μ*m thickness were cut on a cryostat, and mounted on MAS-coated glass slides (Matsunami Glass). Retinal sections were first washed with PBS, blocked for 1-2 h in blocking buffer at RT, incubated with primary antibodies for overnight at 4°C in antibody reaction buffer. The bound antibodies were detected with secondary antibodies after 2 h incubation at 4°C in antibody reaction buffer. The fluorescent samples were counterstained with DAPI. After a final washing in PBS, retinae were embedded in ProLong™ Glass Antifade Mountant (ThermoFisher Scientific).

#### Brain samples

To dissect adult brain, *Igfn1*^*iCre*^; Rosa-LSL-tdTomato adult mice were anesthetized using isoflurane and were transcardially perfused with 4% PFA in PBS. The immediately dissected brains were post-fixed with 4% PFA overnight and later immersed in high sucrose content (30% sucrose in PBS) for 24 to 36 hours. The dissected brains were frozen in OCT compound. Serial coronal sections were cut on a cryostat at a thickness of 30 *μ*m. To visualize tdTomato labelled cells, coronal sections were first incubated with primary antibodies diluted in antibody reaction buffer and were kept for 2 days at 4 °C. Next, after few washes, the sections were incubated in secondary antibody and were also kept overnight at 4 °C. All the fluorescent samples were counterstained with DAPI and mounted on glass slides.

### c. Image acquisition

The stained retinas were imaged using a confocal microscope, Olympus FV 3000, with 60x objective (1.42 NA). The images were acquired at 1024 × 1024 pixel (0.102 *μ*m/pixels). For imaging sections, the thickness of the step size along z-axis was 1 *μ*m. The images were processed and analyzed using Image J (Fiji). For cell density analyses, stained retinas were imaged using 20x objective (1.00 NA). Images were taken from GCL to INL and the thickness of each imaging plane in z stack was 1*μ*m. The z-stacks images were later processed and analyzed using Fiji. For measuring the retina size, DAPI-stained whole mount retina was imaged using an All-in-one Fluorescence Microscope (BZ-X700, Keyence) with an objective lens of CFI 4× Plan Apo Lamda, NA 0.13 (Nikon). For imaging brain sections, Keyence microscope with an objective lens of CFI 10× Plan Apo Lamda, NA 0.45 (Nikon) was used.

### d. Image analysis

The tdTomato cell densities at postnatal stages, P8-P15, were manually counted using cell counter from Fiji on the z-stack confocal images taken from 20x objective. The three nuclear layers were separated using DAPI and ChAT counter staining. To measure cell density in adult samples, open-source code (Auti) was used. All the plots were made in python using matplotlib. To compare the fraction of bipolar and amacrine cells at postnatal stage P12-P15, only those cells which showed distinct INL (bipolar and amacrine) cell morphologies were used. Bipolar cells were distinguished from amacrine cells based on the presence of two distinct processes, dendrites extending towards OPL and axons projecting to the IPL. BC 7 were identified based on axon terminal stratification within sublamina S4 of the IPL. The sublaminar boundaries in IPL were marked by staining SACs by ChAT antibody.

The intensity plot code used to distinguish bipolar cells at P15 was developed in Python using Visual Studio Code. In our analyses, only sparsely labelled bipolar cells which showed clear cell morphologies including apical dendrites and basal axon stratification were used. The two images, representing ChAT and tdTomato channels, were first loaded in color format and then converted to greyscale using OpenCV to standardize intensity values. Greyscale intensity data was extracted and stored as 2D matrices. For each image, the vertical intensity profile was computed by summing pixel intensities along the horizontal axis, followed by normalization. The pixel-to-micrometer conversion factor (0.102) was applied to translate pixel distances into micrometer-scale spatial coordinates. The resulting intensity profiles, expressed in arbitrary units (a.u.) and plotted against depth in micrometers, were visualized to compare signal distribution along the tissue depth.

To measure the total retinal area, whole-retina images stained with DAPI were analyzed in Fiji/ImageJ. The measured area values (µm^2^) were determined and used to estimate retinal size by calculating the equivalent circular diameter. Since, retina does not form a perfect circle, an equivalent diameter was calculated as the diameter of a circle with the measured area. The equivalent diameter (D) was calculated using *D* = 2\*sqrt*{\*frac*{*A*}{\*pi*}}, where A denotes the measured retinal area. The calculated diameters were converted from micrometers to millimeters and used to compare retinal size across genotypes.

To accurately assign anatomical location to the coronal sections, QUINT workflow softwares-QuickNII and VisuAlign (Yates et al. (2019)) registered to Allen Mouse Brain Atlas were used.

### e. Statistical analysis

Normality of the data was assessed Shapiro-Wilk test and variance homogeneity using Levene’s test. As the assumptions of normality and homogeneity of variance were met, parametric tests were used. One way ANOVA and post hoc Tukey test were used to compare mouse body weights and retina size at adult stage among different genotypes (Figure 1B, C), INL and GCL density at postnatal stages (Figure 2C, D). Student’s t-test (pooled) was done to compare cell density between P15 and adult (Figure 3C), and tdTomato and ChAT cell densities in the center and periphery of INL and in the center and periphery of GCL (Figure 3D, Supplementary Figure S1E). All the analysis was done in python by custom made codes.

**Figure 1:**
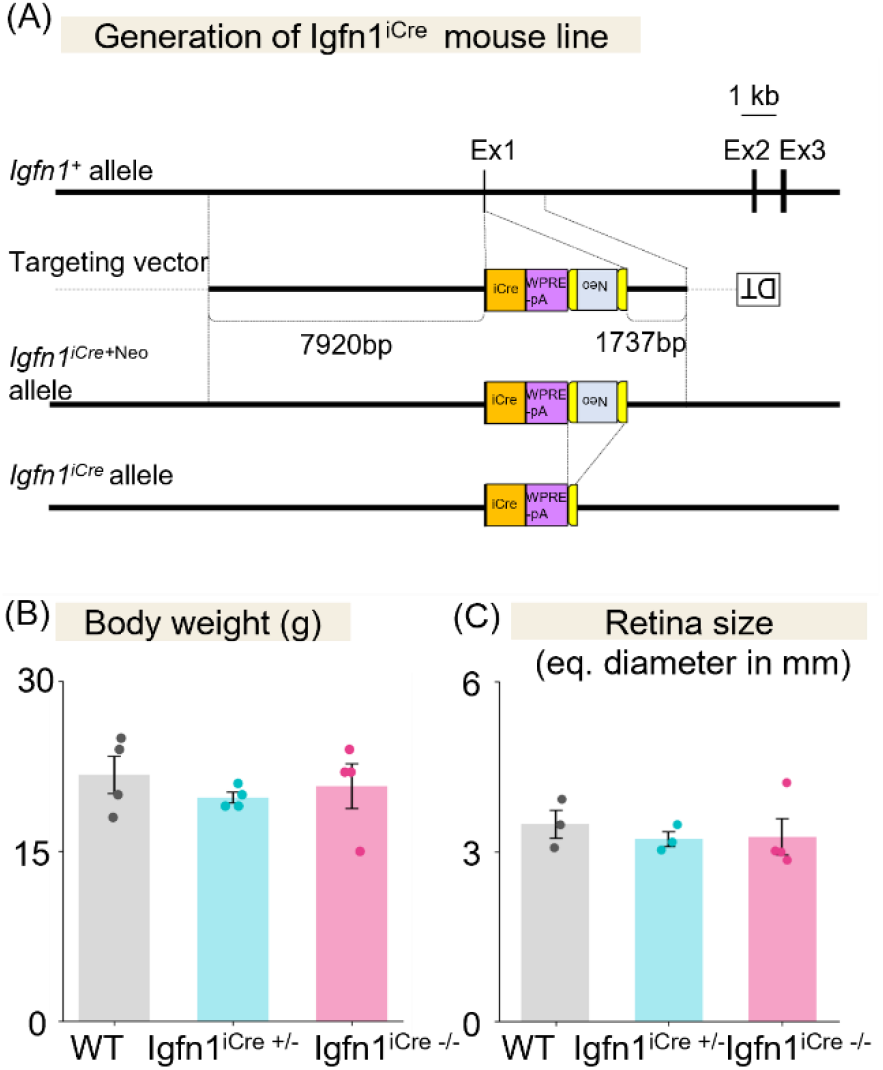
(A) Schematic of the Igfn1 genomic locus. An iCreS/wpre/pA cassette was inserted into exon 1 immediately downstream of the ATG start codon. The approximate lengths of genomic regions flanking the cassette are indicated as homologous arm used for recombination. The selection cassette (Neo) was flanked by FRT sites and excised in the final allele. Exon is abbreviated as Ex. (B) Body weight (in grams) comparison of wild-type (WT), Igfn1^iCre +/-^ (heterozygous), and Igfn1^iCre -/-^ (homozygous) adult male mice. Dots represent individual samples; error bars, mean ± SD (n = 4 adult males). One-way ANOVA with post hoc Tukey-Kramer test (p > 0.05). (C) Retinal size, measured as equivalent diameter (*μ*m), in WT Igfn1^iCre +/-^, and Igfn1^iCre -/-^ adult mice. Dots represent individual samples; error bars, mean ± SD (n = 3 or 4 adult right eyes). One-way ANOVA with post hoc Tukey-Kramer test (p > 0.05).

**Figure 2.**
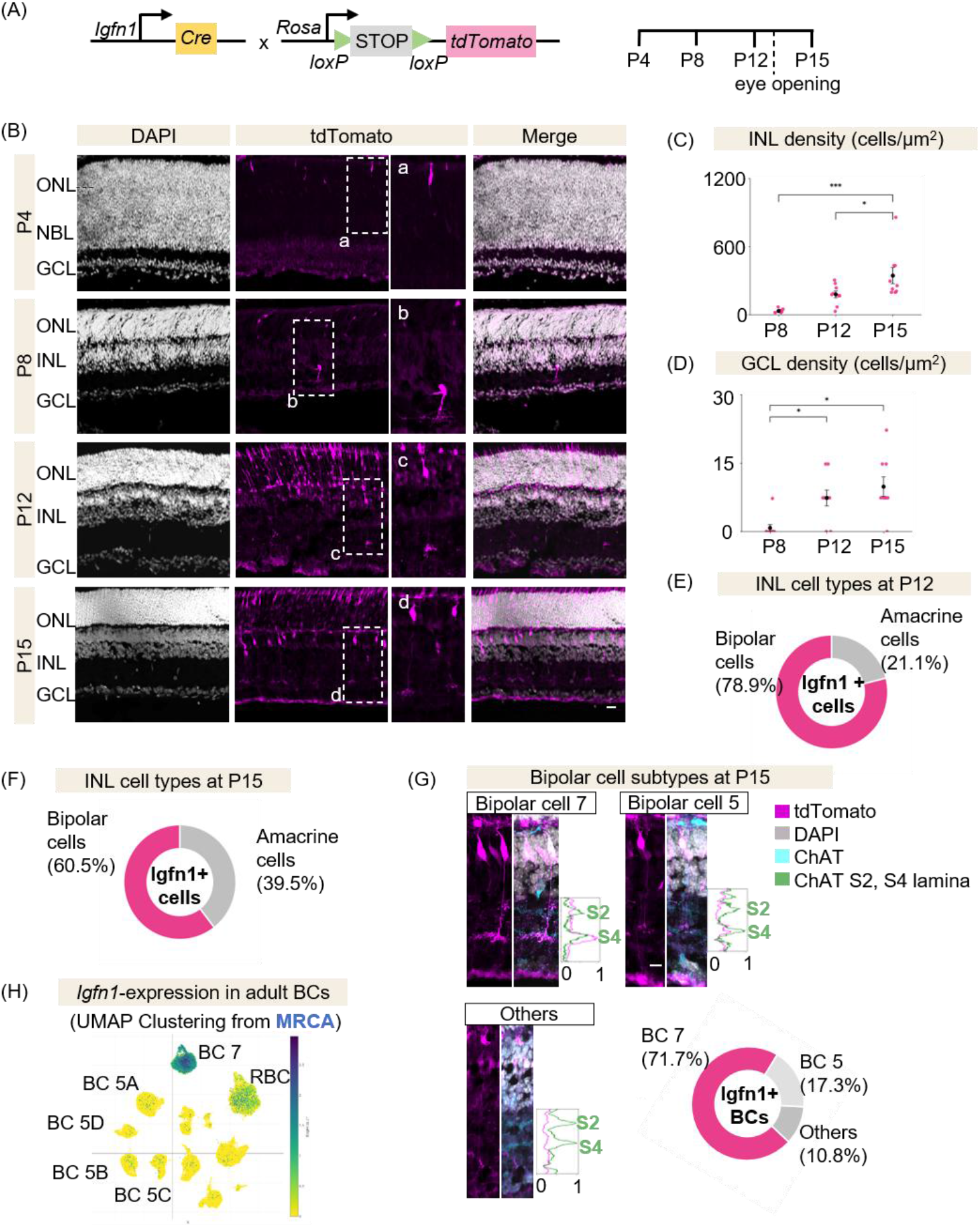
(A) Breeding strategy to generate Igfn1^iCre+/-^; Rosa26-LSL-tdTomato mice+/-mice and dissect retina at postnatal days 4 (P4), P8, P12, P15. (B, inset a-d) Confocal images of retinal sections from Igfn1^iCre +/-^; Rosa26-LSL-tdTomato mice shows tdTomato expression across developmental stages. High magnification of the tdTomato signals in photoreceptor layer in (a), inner nuclear layer (b-d). NBL, neuroblast layer; ONL, outer nuclear layer; INL, inner nuclear layer; GCL, ganglion cell layer. Scale bar, 20 *μ*m. (C) INL cell density (cells/*μ*m2) at P8, P12, and P15. Dots represent individual samples; black dot represents the mean; error bars, mean ± SD (n = 3 images per postnatal stage from 3 wholemount retinas). One-way ANOVA with post hoc Tukey-Kramer test (***p < 0.001, *p < 0.05). (D) GCL cell density (cells/*μ*m2) at P8, P12, and P15. Dots represent individual samples; black dot represents the mean; error bars, mean ± SD (n = 3 images per postnatal stage from 3 wholemount retinas). One-way ANOVA with post hoc Tukey-Kramer test (*p < 0.05). (E, F) Fraction of INL cell types labeled at P12 (n= 57, 3 retinas) and P15 (n= 76, 4 retinas). (G) Confocal images of representative bipolar cell morphologies at P15. tdTomato-labeled axon projections stratify relative to ChAT bands S2 and S4 in the inner plexiform layer (IPL). Scale bar, 10 *μ*m. (H) UMAP of Mouse Retina Cell Atlas (MRCA) single-cell RNA-seq data showing Igfn1 expression across all bipolar cell types.

**Figure 3:**
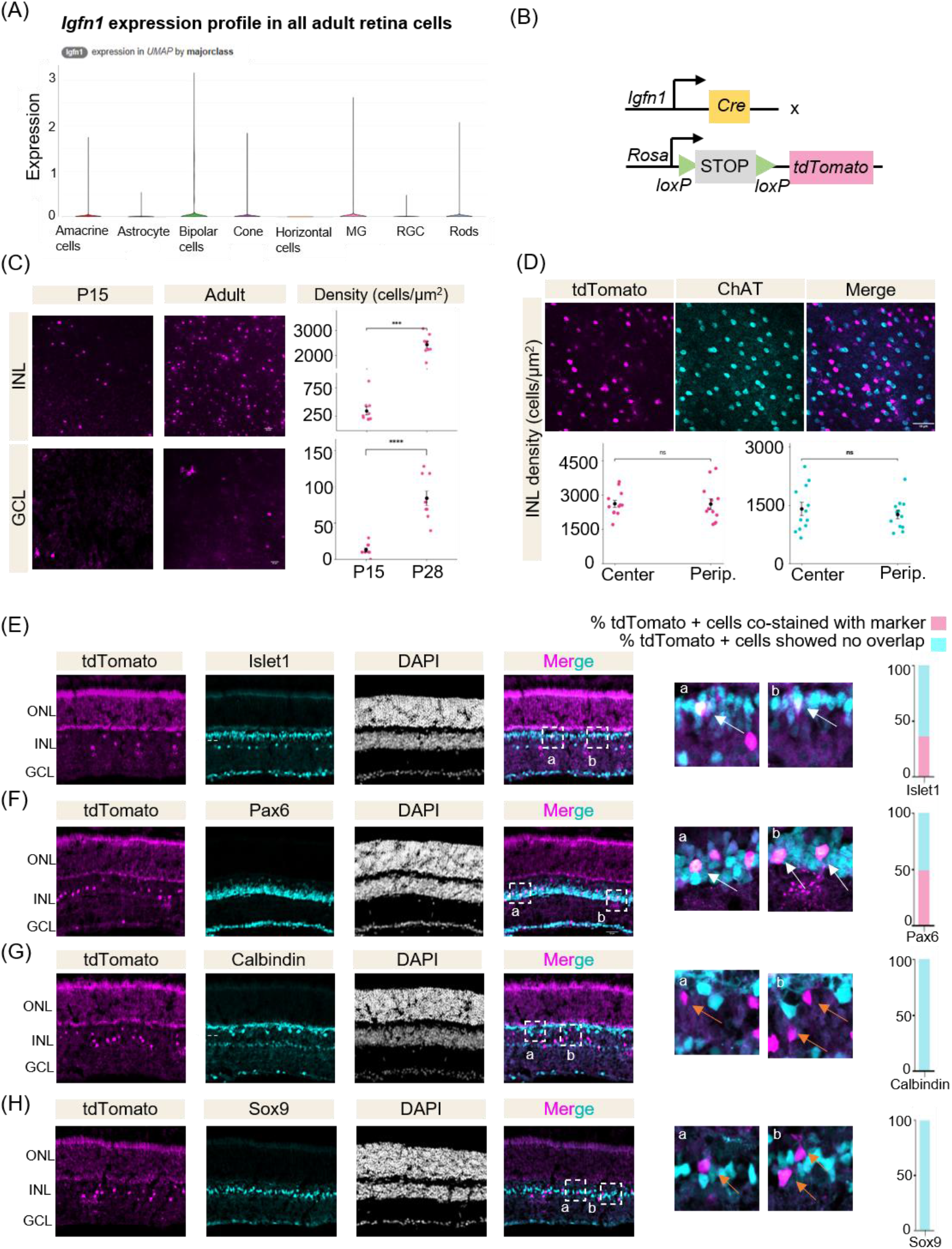
(A) Igfn1 expression across major retinal classes. UMAP from MRCA database. (B) Breeding strategy to generate Igfn1^iCre+/-^; Rosa26-LSL-tdTomato +/-mice and dissect adult retinas. (C) Comparison of INL and GCL tdTomato-positive cell density at P15 and adult stages. Dots represent individual samples; black dot represents the mean; error bars, mean ± SD (n = 3 images per stage from 3 wholemount retinas). For INL density, Student’s t-test (***p < 0.001). For GCL density, Student’s t-test (****p < 0.0001). Scale bar, 20*μ*m. (D) Wholemount image of INL showing ChAT, tdTomato, and merged channels. Graph shows spatial distribution of tdTomato-positive (in magenta) and ChAT-positive cell densities (in cyan) in the INL at central and peripheral regions. Student’s t-test (p > 0.05). Scale bar, 10*μ*m. (E) Islet1 immunostaining in adult retinal sections. Panels I and II show high-magnification images; white arrows indicate colocalization. Islet1^+^/tdTomato^+^ cells in the INL were quantified and their fraction is represented in magenta in the stacked bar graph. (F) Pax6 immunostaining in adult retinal sections. Panels I and II show high-magnification images; white arrows indicate colocalization. Pax6^+^/tdTomato^+^ cells in the INL were quantified and their fraction is represented in magenta in the stacked bar graph. Scale bar, 20*μ*m. (G) Calbindin immunostaining in adult retinal sections. Panels I and II show high-magnification images; yellow arrows indicate no overlap between Calbindin and tdTomato signals. Calbindin^+^/tdTomato^+^ cells in the INL were quantified. (H) Sox9 immunostaining in adult retinal sections. Panels I and II show high-magnification images; yellow arrows indicate no overlap between Sox9 and tdTomato signals. Sox9^+^/tdTomato^+^ cells in the INL were quantified.

## 4. Results

### a. Generation of the *Igfn1*^*iCre*^ recombinase mouse line

Transcriptomic and developmental studies identified *Igfn1* as a BC 7-enriched gene (Shekhar et al. (2016); West et al. (2022)). Thus, to obtain genetic access to subtype 7, we generated a knock-in *Igfn1*^*iCre*^ mouse line, wherein iCre-splice gene cassette was inserted into *Igfn1* gene locus downstream of endogenous, cell-type-specific promoter (Figure 1A). Both, heterozygous and homozygous *Igfn1*^*iCre*^ mice were viable, fertile, and displayed normal body weight, compared to the wild type controls (Figure 1B). The overall retina size of the heterozygotes was indistinguishable from wild-type controls (Figure 1C). This indicates that there are no developmental or retina structural abnormalities in the or homozygous transgenic mice. Thus, we proceeded with *Igfn1*^*iCre*^ as a driver line to further study the type-7 bipolar cell.

### b. Developmental onset and cell type diversity of *Igfn1*^*iCre*^ expression in the retina

The developmental onset of *Igfn1* expression and its cell type specificity was characterized by crossing the *Igfn1*^*iCre*^ mice with a Cre-dependent tdTomato reporter mice and analyzed in retinal sections and flat mounts from P4 to P15 (Figure 2A). To classify the tdTomato-expressing cells we first determined their soma position within the neuroblast layer (NBL), outer nuclear layer (ONL), inner nuclear layer (INL), or ganglion cell layer (GCL). We then identified the labelled cells by the morphology of their dendritic and axonal projections. The neurons located in the ONL are considered photoreceptors. The cells in INL and that extends processes into the IPL were categorized as bipolar or amacrine cells. These neurons can be segregated into bipolar cells and amacrine cells by presence or absence of stout dendrites extending toward the Outer Plexiform Layer (OPL) respectively (Zhu et al. (2014)). Additional INL neurons include horizontal cells which extends neurites only to OPL and glia cells which spans the retina from ONL to GCL. The GCL consists of retinal ganglion cells and displaced amacrine cells which can be identified based on the presence or absence, respectively, of tdTomato signal in an axon coursing toward the optic nerve head or within the optic nerve.

At P4 stage retinal cross sections, we observed tdTomato expression was restricted to the outer parts of NBL, corresponding to the presumptive outer nuclear layer (ONL) as defined in previous studies (Postel et al. (2013)), and the labelled cells displayed morphologies similar to photoreceptors (Figure 2B, inset a). By P8, a sparse population of tdTomato-positive cells were visible in the INL, however, some of these cells did not display clear bipolar-like features. Some labelled cells displayed two neurites towards OPL and IPL whereas others extended only one neurite projection to the IPL (Figure 2B, inset b). This may suggest labelling of non-BCs types at this stage. Thus, to evaluate whether these were amacrine cells, we immunostained for Pax6, a pan-amacrine marker expressed in both conventional and displaced amacrine cells (Kim et al. (2017)). We observed a heterogeneous population as some tdTomato-positive cells were Pax6-positive, while others were Pax6-negative (Figure S1A). The two groups could be differed by soma position and presence or absence of apical dendrites as Pax6-negative soma lay closer to the OPL, and shows presence of an apical and basal dendrite, and perhaps represent bipolar cells, whereas Pax6-positive soma localized at lower INL and projects its neurite to IPL only, consistent with amacrine cells. Further, at P12, retinal sections showed tdTomato-positive cells with hallmark bipolar morphology, including clear apical and basal processes (Figure 2B, inset c). The number of tdTomato-expressing cells in the INL increased substantially by P15, forming a heterogeneous population with both bipolar and amacrine cell morphologies (Figure 2B, inset d). Whole-mount quantification also showed a progressive increase in INL tdTomato cell density from P8 to P15 (Figure 2C).

We classified tdTomato-expressing cell types at P12–P15 by examining retina sections. Applying the criteria above, approximately 80% of labelled INL cells at P12 and 60% at P15 were bipolar cells, and the remaining cells identified as amacrine cells (Figure 2E–F). In the absence of a distinctive soma histochemical marker for bipolar cell subtypes, we relied on documented literature of axon stratification as the primary criterion to further identify bipolar cell subtype-7 (West and Cepko (2022); Shekhar et al. (2016); Euler et al. (2014)). At P15, about 71% bipolar cells stratified in IPL sublamina S4, immediately beneath the ON ChAT band marked by starburst amacrine cell processes, which is a classical laminar position of BC 7 (Figure 2G) (Matsumoto et al. (2021); Shekhar et al. (2016); Wässle et al. (2009); Hall et al. (2019)). A smaller fraction, about 17%, projected axons on the ON ChAT band, consistent with BC 5 identity (Sharpe et al. (2022); Hellmer et al. (2016); Hall et al. (2019)), and the remainder stratified much deeper in the IPL, resembling rod BCs or other subtypes (Hall et al. (2019); Ghosh et al. (2004); Huh et al. (2015)(Figure 2G)). Interestingly, we also observed that tdTomato-positive bipolar cells at P12, stratified in the sublamina S4, identical to subtype 7 morphologies at P15 (Figure S1B-C). This observation suggests that that *Igfn1*^*iCre*^ provides limited access to the subtype before P12–P15 and potential access at around P15.

As a complementary analysis to compare *Igfn1*^*iCre*^ dependent reporter labelling in bipolar cells with the endogenous *Igfn1* transcript distribution, we checked *Igfn1* expression in the recently available and publicly accessible Mouse Retina Cell Atlas (MRCA) of the adult wild-type retina (Li et al. (2024)). We found within the bipolar cell class, *Igfn1* transcripts were enriched in BC 7, but also expressed in other subtypes including BC 5 clusters, rod BCs (Figure 2H), consistent with our experimental observations.

To further characterize the *Igfn1*^*iCre*^ transgenic mouse line, we examined bipolar cell stratification in homozygous mice crossed with the Rosa-LSL-tdTomato reporter at P15. In the retina, DAPI staining revealed intact lamination of all the nuclear layers, with no apparent structural abnormalities (Figure S2A). Within the INL, tdTomato labeled cells comprised a mixed neuronal population, with approximately 60% classified as bipolar cells and the remainder as amacrine cells, similar to the heterozygous condition (Figure S2B). We further analyzed axon terminal stratification and found approximately 80% of the labeled bipolar cells projected to sublamina S4, consistent with BC 7 identity, while additional cells displayed stratification patterns corresponding to other bipolar subtypes (Figure S2C). Importantly, the laminar positioning of BC 7 axon terminals within S4 were indistinguishable between homozygous and heterozygous samples. These findings suggest that disruption of exon 1 in the *Igfn1* knock-in allele does not produce gross defects in bipolar cell stratification at this developmental stage. Several explanations remain possible. Either *Igfn1* may not play a major structural role in axon targeting of BC 7 cells or disruption of exon 1 may not eliminate critical functional domains of the protein such as immunoglobulin-like or fibronectin motifs, thereby preserving bipolar structure. Additional molecular or functional analyses will be required to clarify the contribution of *Igfn1* to bipolar cell development.

In addition to BCs, we observed tdTomato-positive amacrine cells with distinct stratification profiles in the IPL (Figure S1D), indicating that *Igfn1*^*iCre*^ also labels a few amacrine cell populations. In the GCL, labelled tdTomato-positive cells were sparse with wide dendritic arbors in the IPL, lacked axons coursing along the vitreal surface, and almost no signal in the optic nerve. The labelled GCL cell density increased significantly from P8 to P12 and did not change further from P12 to P15 (Figure 2 D). Across all stages, labelling remained more prominent in the INL than in the GCL.

Together, these findings show that *Igfn1*^*iCre*^ drives tdTomato expression in a developmental pattern in diverse retinal cells. Our findings shows that reporter signal first appears in the outer retina labelling photoreceptors and later extends to bipolar and amacrine cells in the inner retina. Although bipolar cell subtype genesis peaks around P3–P4, type 7 labelling does not become prominent until P12–15, which is coincident with bipolar cell maturation period. Overall, by P12–P15, the transgenic line can provide partially selective access to BC 7, a functionally significant subtype implicated in direction-selective retinal circuit.

### c. Spatial and histochemical characterization of *Igfn1*^*iCre*^ labelled cells in the adult retina

The single cell RNA-seq data, as previously mentioned, identified *Igfn1* as an adult-enriched marker of BC 7 (Shekhar et al. (2016)). Thus, to validate our experimental findings in additional non-bipolar cell types, we examined *Igfn1* expression across all retinal populations using the MRCA atlas. As shown in (Figure 3 A), *Igfn1* expression was enriched in BC 7 but not exclusive to this subtype as comparatively lower transcripts were also detected in photoreceptors, amacrine cells, and glia cells, whereas minimal to no expression in ganglion cells and horizontal cells. Further, we found the *Igfn1* mRNA were distributed across approximately 40 amacrine cell clusters (Figure S3 A) corresponding to various transmitter categories. Together, these data provide a plausible molecular explanation for the broader cellular labelling observed in our experiments.

We therefore asked whether the *Igfn1*^*iCre*^ driver can potentially provide robust access to adult bipolar cell type, and how labelling is distributed between INL cell populations at this stage. To address this, we applied the previously mentioned strategy using tdTomato reporter mice and imaged retinal flat mounts and vertical sections at adult stage (Figure 3B). In retinal whole mounts, to segregate nuclear layers, we used ChAT immunostaining to mark OFF SAC soma in the INL and ON SAC soma in the GCL as anatomical landmarks (Figure 3C–D). We observed that tdTomato-positive cell density in both the GCL and INL was significantly higher in adult retinas compared to P15, indicating a developmental increase in labelled cells (Figure 3C). To assess whether labelling varied across retinal regions, we quantified tdTomato-positive densities in central and peripheral regions in the inner retina. In both the nuclear layers, tdTomato-positive cell densities showed no significant difference between center and periphery, suggesting uniform cell distribution in the layers (Figure 3D, Figure S1E). We also characterized ChAT-positive SACs densities as a positive control to ensure uniform distribution (Figure 3D, Figure S1E).

As high cell densities precluded reliable morphology-based classification, we used immunohistochemistry in retinal cross-sections to identify tdTomato labelled cells. To classify labelled neurons, we used well documented markers, prominently expressed in the cell body. We first stained for Islet1, a marker of ON bipolar cells, noting its expression in subsets of SACs and retinal ganglion cells (Elshatory et al. (2007)). In the INL, Islet1-positive nuclei appeared in two discrete bands (marked by a white line in the Figure 3E, column 2), an upper band corresponding to ON bipolar cells and a lower band representing SACs. Approximately 40% of tdTomato-expressing INL cells colocalized with Islet1 (see insets a& b of Figure 3E), and many of these double-positive cells displayed vertically oriented processes to OPL consistent with bipolar morphology. Further, to evaluate amacrine cell labelling, we immunostained for Pax6 (Kim et al. (2017)) and found approximately 50% of tdTomato-expressing INL cells colocalized with Pax6 (see insets a& b of Figure 3F), indicating that a substantial portion of the labelled population comprises amacrine cells. Finally, we stained for Calbindin, a horizontal cell marker (Haverkamp and Wässle (2000)). We found no overlap between calbindin positive horizontal cells located above the white line in the (Figure 3G, column 2) and tdTomato-expressing cells (see insets a & b of Figure 3G), indicating that *Igfn1*^*iCre*^ does not label horizontal cells. Further, in the GCL, most tdTomato positive cells co-stained with Pax6 and none with Calbindin, and these cells showed no clear signal in axons projecting to optic nerve head, thereby suggesting displaced amacrine cell identity (Figure S1F). At last, we checked whether *Igfn1*^*iCre*^ targets non-neuronal cells by co-staining with Sox9, which marks Müller glia in the INL and astrocytes in the GCL (Yamamoto et al. (2020); Fischer et al. (2010)). Sox9-positive nuclei were evident in both layers, but none overlapped with tdTomato-expressing cells (see insets a & b of Figure 3H), indicating that *Igfn1*^*iCre*^ does not label retinal glial cells. Thus, *Igfn1*^*iCre*^ expression in the adult retina is confined to neuronal populations.

Together, these data show that in the adult retina, there is a drastic increase in the tdTomato-expressing cell densities, and the labelled INL population consists of ON bipolar and amacrine neurons, with no horizontal or glial cell labelling. These adult histological findings are largely consistent with the MRCA atlas results, with the exception of glial cells. One possible explanation is *Igfn1* can harbor multiple potential translation initiation sites, that could be regulated by other promoter/enhancers, resulting in expression of different protein isoforms, as reported previously in non-neuronal study (Li et al. (2017)). As the iCre-cassette insertion was in exon 1, the recombination may have surprisingly happened only in neurons in the retina.

### d. *Igfn1*^*iCre*^ Expression Across Brain Regions in the Adult Mouse

The *Igfn1*^*iCre*^ mouse line can be utilized for studies beyond the retina. Thus, to determine whether *Igfn1*^*iCre*^ drives reporter expression in central visual areas and other brain regions, we examined *Igfn1*^*iCre*^; Rosa-LSL-tdTomato adult brain coronal sections and high-resolution imaging (Figure 4A). The images sections were processed using QUINT workflow (Yates et al. (2019)) registered to Allen Mouse Brain Atlas to accurately annotate brain regions (Figure S4A– B).

**Figure 4.**
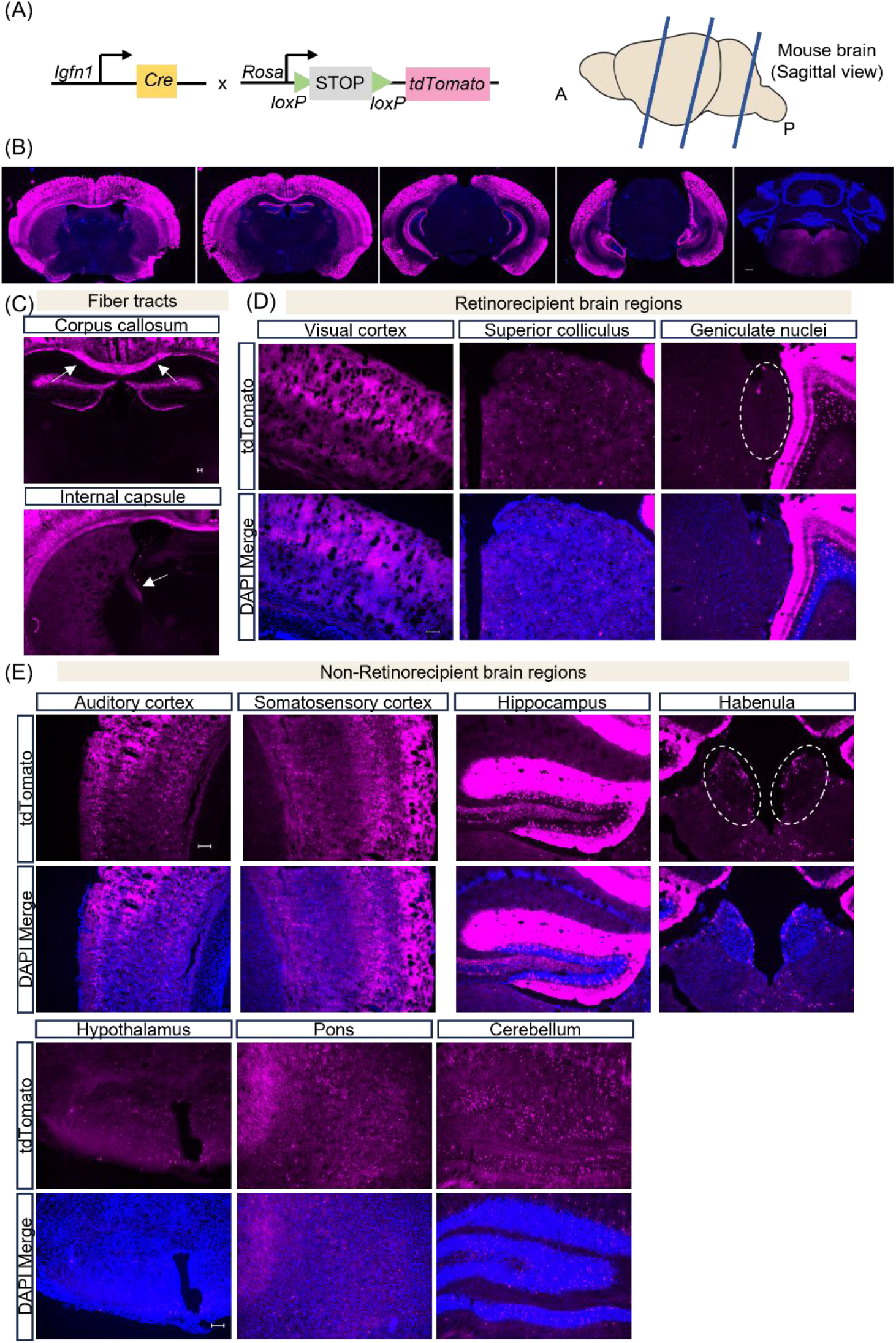
(A) Left, Breeding strategy to generate *Igfn1* ^*iCre+/-*^; Rosa26-LSL-tdTomato^+/-^ mice and dissect adult brains. Right side is schematic of the adult mouse brain (sagittal view) and blue lines indicate the angle of sectioning for coronal slices. (B) Stitched fluorescent image of tdTomato reporter expression in a coronal brain section. Scale bar, 600 *μ*m. (C) High-magnification fluorescent images showing tdTomato expression in brain fiber tracts. Scale bar, 100 *μ*m. (D–E) High-magnification fluorescent images showing tdTomato expression in representative brain regions. Scale bar, 100 *μ*m.

We observed dense tdTomato fluorescence in the anterior forebrain brain regions, including the cortex and hippocampus and much sparser expression in posterior midbrain and hindbrain areas with an exception of cerebellum (Figure 4B). Additionally, tdTomato labelling was also present in fiber tracts including the corpus callosum, internal capsule suggesting labelled population includes long-range projection neurons (Figure 4C). We next imaged the central visual areas including visual cortex, superior colliculus (SC), and geniculate nucleus. The visual cortex displayed tdTomato labelling across multiple cortical layers, in contrast to the superficial SC and geniculate nucleus, where tdTomato signal was sparse and did not show any laminar organization or was absent (Figure 4D).

In the non-visual cortical areas, tdTomato expression spanned from anterior motor cortex to posterior retrosplenial cortex. High magnification images of cortical regions including primary somatosensory and auditory cortices show higher density of labelled soma concentrated in several layers, including deep layers 5 and 6 (Figure 4E). In subcortical forebrain regions, reporter expression was prominent in the dentate gyrus, with tdTomato-positive cells spanning both the entire structure (Figure 4E). We also noted sparse labelling in the medial habenula. In the hypothalamus, tdTomato-positive cells localized to scattered ventral areas, including the periventricular hypothalamus and arcuate nucleus (Figure 4E). In the hindbrain areas, sparse labelling was detected in the pontine and medullary reticular formations (Figure 4E) whereas comparatively denser signals were present across cerebellum lobules.

Overall, these findings characterize the *Igfn1*^*iCre*^ mouse line, showing its widespread labelling across the adult brain, with region-dependent differences in labelling density.

## 5. Discussion

### a. Significance of the postnatal BC 7 accessibility

Our characterization results reveal a clear temporal sequence of *Igfn1*^*iCre*^ expression in relation to BC 7. Sparse labelling was detected in the INL as early as P8, increased substantially by P12–P15, and expanded further in adult. This developmental trajectory aligns with the period of bipolar cell maturation which includes axon stratification, synapse formation, rather than with early cell fate specification (West and Cepko (2022)). Thus, it suggests *Igfn1* may have a structural role during BC 7 differentiation.

The postnatal window around eye opening represents an important stage for experimental access to BC 7. This period coincides with multiple developmental programs, including spontaneous retinal activity (Stage II–III retinal waves) (Voufo et al. (2023)) and the formation of synaptic connections between SACs and output ganglion cells (Yonehara et al. (2011)), which are key circuit-level events underlying direction selectivity. We speculate this postnatal period can provide a starting framework for future studies of development of feature selectivity in BC 7. These studies can potentially offer interesting insights into the formation of neural circuits in the mammalian nervous system.

### b. BC 7 access window and limitations

For studies requiring BC 7-enriched access, P15 represents an “accessible spot”. At this time point, a large fraction of labelled bipolar interneurons stratifies in S4 beneath the ON ChAT band, consistent with BC 7 identity. By contrast, in the adult retina the transgenic mouse line’s utility for BC 7-specific manipulations are limited due to the slightly higher amacrine labelling. To achieve more selective type-7 access in adults, we suggest intersectional strategies. One viable route is to combine *Igfn1*^*iCre*^ with an ON-bipolar enriched Flp driver (e.g., Grm6–Flp) so that only neurons satisfying both conditions (Cre and Flp) are labelled, thereby enriching BC 7 while excluding most amacrines and non-target BCs. Alternatively, generation of an inducible *Igfn1*^*CreER*^ mouse line, along with administration of tamoxifen between P12 and P15, could restrict recombination in BC 7.

### c. Comparison of reporter labeling and ISH data in the adult brain

Our characterization results using a reporter mice in the adult brain broadly matches to the *Igfn1* expression pattern reported in the Allen Brain Atlas, particularly in anterior forebrain brain regions, including cortical and hippocampus (Lein et al. (2007); Allen Institute for Brain Science (2004a,b)). However, we observed tdTomato labeling in the hindbrain region cerebellum which is not indicated by the in-situ hybridization (ISH) dataset. Although the role of *Igfn1* in the nervous system has not been established, the observed cerebellar reporter signal could be due to the following factors. As *Igfn1*^*iCre*^ induces permanent tdTomato recombination, transient *Igfn1* expression during early postnatal stages could result in reporter labelling even if adult expression is low or undetectable by ISH. In addition, Cre-dependent reporter detection is more sensitive than ISH and can reveal low-level gene expression, that may fall below ISH detection threshold (Heffner et al. (2012)). Moreover, the ISH probe design may not fully capture splice variant diversity of *Igfn1*, leading to undetectable transcript isoforms in certain brain regions.

## 6. Conclusion

This study presents the first genetic line, *Igfn1*^*iCre*^, designed to access a bipolar cell subtype based on cell type enriched molecular marker. Through morphological and histological analyses, we characterize the developmental profile of type-7 marker gene, *Igfn1*, which is first expressed in the outer retina and then expands into the inner nuclear layer to bipolar and amacrine cells (Figure S3 B). Importantly, BC 7-enriched labelling emerges around P12–P15, marked with axonal stratification in S4 plexiform layer. This represents the first case in which a bipolar subtype defined by an adult scRNA-seq marker provides genetic access to BC 7 during a developmental window near eye opening. The widespread expression patterns in the adult retina and brain highlight the need for additional strategies to access target neurons.

## CRediT authorship contribution statement

**Shambhavi Chaturvedi:** Conceptualization, Methodology, Formal analysis, Visualization, Validation, Writing – original draft. **Haruka Yamamoto:** Conceptualization, Supervision, Writing – review and editing. **Akihiro Matsumoto:** Supervision, Writing – review and editing. **Manabu Abe:** Methodology, Funding acquisition, Writing – review and editing. **Toshikuni Sasaoka:** Methodology, Funding acquisition, Writing – review and editing. **Keisuke Yonehara:** Conceptualization, Supervision, Funding acquisition, Project administration, Writing – review and editing.

## Acknowledgements

We thank all Yonehara lab members for guidance and discussion. We are grateful for the technical assistance provided by Yonehara lab technicians including Makoto Kiso, Madoka Hagihara and Chiaki Sakiyama. S. Chaturvedi was supported by the Japanese government Monbukagakusho (MEXT) scholarship. This work was supported by funding from KAKENHI (20K23377; 22K21353; 23H04241; 24H02311), the Collaborative Research Project (#24013, #25015) of Brain Research Institute, Niigata University, the Global Collaborative Research Project (G2905, G3005, G202101) of Brain Research Institute, Niigata University, Chugai Foundation for Innovative Drug Discovery Science, Daiichi Sankyo Foundation of Life Science, Mochida Memorial Foundation for Medical and Pharmaceutical Research, Mitsubishi Foundation, Toray Science Foundation, and Naito Foundation to K.Yonehara.

## Data Availability

The data supporting the findings of this study are available from the corresponding author upon request.

## References

Allen Institute for Brain Science, 2004a. Allen mouse brain atlas. Dataset. Available from: https://mouse.brain-map.org/.

Allen Institute for Brain Science, 2004b. Igfn1 – experiment detail. allen mouse brain atlas. Available from: https://mouse.brain-map.org/gene/show/86416.

Auti, G., Image Analysis. https://github.com/gunjansauti/ImageAnalysis.

Elshatory, Y., Deng, M., Xie, X., Gan, L., 2007. Expression of the LIM-homeodomain protein Isl1 in the developing and mature mouse retina. J of Comparative Neurology 503, 182–197. URL: https://onlinelibrary.wiley.com/doi/10.1002/cne.21390, doi:10.1002/cne.21390.

Euler, T., Haverkamp, S., Schubert, T., Baden, T., 2014. Retinal bipolar cells: elementary building blocks of vision. Nat Rev Neurosci 15, 507–519. URL: https://www.nature.com/articles/nrn3783, doi:10.1038/nrn3783.

Fischer, A.J., Zelinka, C., Scott, M.A., 2010. Heterogeneity of Glia in the Retina and Optic Nerve of Birds and Mammals. PLoS ONE 5, e10774. URL: https://dx.plos.org/10.1371/journal.pone.0010774, doi:10.1371/journal.pone.0010774.

Franke, K., Berens, P., Schubert, T., Bethge, M., Euler, T., Baden, T., 2017. Inhibition decorrelates visual feature representations in the inner retina. Nature 542, 439–444. URL: https://www.nature.com/articles/nature21394, doi:10.1038/nature21394.

Ghosh, K.K., Bujan, S., Haverkamp, S., Feigenspan, A., Wässle, H., 2004. Types of bipolar cells in the mouse retina. J of Comparative Neurology 469, 70–82. URL: https://onlinelibrary.wiley.com/doi/10.1002/cne.10985, doi:10.1002/cne.10985.

Goetz, J., Jessen, Z.F., Jacobi, A., Mani, A., Cooler, S., Greer, D., Kadri, S., Segal, J., Shekhar, K., Sanes, J.R., Schwartz, G.W., 2022. Unified classification of mouse retinal ganglion cells using function, morphology, and gene expression. Cell Reports 40, 111040. URL: https://linkinghub.elsevier.com/retrieve/pii/S2211124722008348, doi:10.1016/j.celrep.2022.111040.

Hall, L.M., Hellmer, C.B., Koehler, C.C., Ichinose, T., 2019. Bipolar Cell Type-Specific Expression and Conductance of Alpha-7 Nicotinic Acetylcholine Receptors in the Mouse Retina. Invest. Ophthalmol. Vis. Sci. 60, 1353. URL: http://iovs.arvojournals.org/article.aspx?doi=10.1167/iovs.18-25753, doi:10.1167/iovs.18-25753.

Haverkamp, S., Wässle, H., 2000. Immunocytochemical analysis of the mouse retina. J. Comp. Neurol. 424, 1–23. URL: https://onlinelibrary.wiley.com/doi/10.1002/1096-9861(20000814)424:1<1::AID-CNE1>3.0.CO;2-V, doi:10.1002/1096-9861(20000814)424:1<1::AID-CNE1>3.0.CO;2-V.

Heffner, C.S., Herbert Pratt, C., Babiuk, R.P., Sharma, Y., Rockwood, S.F., Donahue, L.R., Eppig, J.T., Murray, S.A., 2012. Supporting conditional mouse mutagenesis with a comprehensive cre characterization resource. Nat Commun 3, 1218. URL: https://www.nature.com/articles/ncomms2186, doi:10.1038/ncomms2186.

Hellmer, C., Zhou, Y., Fyk-Kolodziej, B., Hu, Z., Ichinose, T., 2016. Morphological and physiological analysis of type-5 and other bipolar cells in the Mouse Retina. Neuroscience 315, 246–258. URL: https://linkinghub.elsevier.com/retrieve/pii/S0306452215010994, doi:10.1016/j.neuroscience.2015.12.016.

Hsiang, J.C., Shen, N., Soto, F., Kerschensteiner, D., 2024. Distributed feature representations of natural stimuli across parallel retinal pathways. Nat Commun 15, 1920. URL: https://www.nature.com/articles/s41467-024-46348-y, doi:10.1038/s41467-024-46348-y.

Huang, L., Max, M., Margolskee, R.F., Su, H., Masland, R.H., Euler, T., 2003. G protein subunit Gγ13 is coexpressed with Gαo, Gβ3, and Gβ4 in retinal ON bipolar cells. J of Comparative Neurology 455, 1–10. URL: https://onlinelibrary.wiley.com/doi/10.1002/cne.10396, doi:10.1002/cne.10396.

Huh, Y.J., Choi, J.S., Jeon, C.J., 2015. Localization of Rod Bipolar Cells in the Mammalian Retina Using an Antibody Against the α1c L-type Ca2+ Channel. Acta Histochemica et Cytochemica 48, 47–52. URL: https://www.jstage.jst.go.jp/article/ahc/48/2/48_14049/_article, doi:10.1267/ahc.14049.

Inoue, R., Abdou, K., Hayashi-Tanaka, A., Muramatsu, S.i., Mino, K., Inokuchi, K., Mori, H., 2018. Glucocorticoid receptor-mediated amygdalar metaplasticity underlies adaptive modulation of fear memory by stress. eLife 7, e34135. URL: https://elifesciences.org/articles/34135, doi:10.7554/eLife.34135.

Kim, Y., Lim, S., Ha, T., Song, Y.H., Sohn, Y.I., Park, D.J., Paik, S.S., Kim-Kaneyama, J.r., Song, M.R., Leung, A., Levine, E.M., Kim, I.B., Goo, Y.S., Lee, S.H., Kang, K.H., Kim, J.W., 2017. The LIM protein complex establishes a retinal circuitry of visual adaptation by regulating Pax6 α-enhancer activity. eLife 6, e21303. URL: https://elifesciences.org/articles/21303, doi:10.7554/eLife.21303.

Lein, E.S., Hawrylycz, M.J., Ao, N., Ayres, M., Bensinger, A., Bernard, A., Boe, A.F., Boguski, M.S., Brockway, K.S., Byrnes, E.J., et al., 2007. Genome-wide atlas of gene expression in the adult mouse brain. Nature 445, 168–176.

Li, J., Choi, J., Cheng, X., Ma, J., Pema, S., Sanes, J.R., Mardon, G., Frankfort, B.J., Tran, N.M., Li, Y., Chen, R., 2024. Comprehensive single-cell atlas of the mouse retina. iScience 27, 109916. URL: https://linkinghub.elsevier.com/retrieve/pii/S2589004224011386, doi:10.1016/j.isci.2024.109916.

Li, X., Baker, J., Cracknell, T., Haynes, A.R., Blanco, G., 2017. IGFN1_v1 is required for myoblast fusion and differentiation. PLoS ONE 12, e0180217. URL: https://dx.plos.org/10.1371/journal.pone.0180217, doi:10.1371/journal.pone.0180217.

Matsumoto, A., Agbariah, W., Nolte, S.S., Andrawos, R., Levi, H., Sabbah, S., Yonehara, K., 2021. Direction selectivity in retinal bipolar cell axon terminals. Neuron 109, 2928–2942.e8. URL: https://linkinghub.elsevier.com/retrieve/pii/S0896627321005183, doi:10.1016/j.neuron.2021.07.008.

Postel, K., Bellmann, J., Splith, V., Ader, M., 2013. Analysis of cell surface markers specific for transplantable rod photoreceptors. Mol Vis 19, 2058–2067.

Sharpe, Z.J., Shehu, A., Ichinose, T., 2022. Asymmetric Distributions of Achromatic Bipolar Cells in the Mouse Retina. Front. Neuroanat. 15, 786142. URL: https://www.frontiersin.org/articles/10.3389/fnana.2021.786142/full, doi:10.3389/fnana.2021.786142.

Shekhar, K., Lapan, S.W., Whitney, I.E., Tran, N.M., Macosko, E.Z., Kowalczyk, M., Adiconis, X., Levin, J.Z., Nemesh, J., Goldman, M., McCarroll, S.A., Cepko, C.L., Regev, A., Sanes, J.R., 2016. Comprehensive Classification of Retinal Bipolar Neurons by Single-Cell Transcriptomics. Cell 166, 1308–1323.e30. URL: https://linkinghub.elsevier.com/retrieve/pii/S0092867416310078, doi:10.1016/j.cell.2016.07.054.

Stabio, M.E., Sabbah, S., Quattrochi, L.E., Ilardi, M.C., Fogerson, P.M., Leyrer, M.L., Kim, M.T., Kim, I., Schiel, M., Renna, J.M., Briggman, K.L., Berson, D.M., 2018. The M5 Cell: A Color-Opponent Intrinsically Photosensitive Retinal Ganglion Cell. Neuron 97, 150–163.e4. URL: https://linkinghub.elsevier.com/retrieve/pii/S0896627317310838, doi:10.1016/j.neuron.2017.11.030.

Ueno, A., Omori, Y., Sugita, Y., Watanabe, S., Chaya, T., Kozuka, T., Kon, T., Yoshida, S., Matsushita, K., Kuwahara, R., Kajimura, N., Okada, Y., Furukawa, T., 2018. Lrit1, a Retinal Transmembrane Protein, Regulates Selective Synapse Formation in Cone Photoreceptor Cells and Visual Acuity. Cell Reports 22, 3548–3561. URL: https://linkinghub.elsevier.com/retrieve/pii/S2211124718303218, doi:10.1016/j.celrep.2018.03.007.

Voufo, C., Chen, A.Q., Smith, B.E., Yan, R., Feller, M.B., Tiriac, A., 2023. Circuit mechanisms underlying embryonic retinal waves. eLife 12, e81983. URL: https://elifesciences.org/articles/81983, doi:10.7554/eLife.81983.

Wässle, H., Puller, C., Müller, F., Haverkamp, S., 2009. Cone Contacts, Mosaics, and Territories of Bipolar Cells in the Mouse Retina. J. Neurosci. 29, 106–117. URL: https://www.jneurosci.org/lookup/doi/10.1523/JNEUROSCI.4442-08.2009, doi:10.1523/JNEUROSCI.4442-08.2009.

West, E.R., Cepko, C.L., 2022. Development and diversification of bipolar interneurons in the mammalian retina. Developmental Biology 481, 30–42. URL:https://linkinghub.elsevier.com/retrieve/pii/S0012160621002128, doi:10.1016/j.ydbio.2021.09.005.

West, E.R., Lapan, S.W., Lee, C., Kajderowicz, K.M., Li, X., Cepko, C.L., 2022. Spatiotemporal patterns of neuronal subtype genesis suggest hierarchical development of retinal diversity. Cell Reports 38, 110191. URL: https://linkinghub.elsevier.com/retrieve/pii/S2211124721016922, doi:10.1016/j.celrep.2021.110191.

Yamamoto, H., Kon, T., Omori, Y., Furukawa, T., 2020. Functional and Evolutionary Diversification of Otx2 and Crx in Vertebrate Retinal Photoreceptor and Bipolar Cell Development. Cell Reports 30, 658–671.e5. URL: https://linkinghub.elsevier.com/retrieve/pii/S2211124719317292, doi:10.1016/j.celrep.2019.12.072.

Yan, W., Laboulaye, M.A., Tran, N.M., Whitney, I.E., Benhar, I., Sanes, J.R., 2020. Mouse Retinal Cell Atlas: Molecular Identification of over Sixty Amacrine Cell Types. J. Neurosci. 40, 5177–5195. URL: https://www.jneurosci.org/lookup/doi/10.1523/JNEUROSCI.0471-20.2020, doi:10.1523/JNEUROSCI.0471-20.2020.

Yates, S.C., Groeneboom, N.E., Coello, C., Lichtenthaler, S.F., Kuhn, P.H., Demuth, H.U., Hartlage-Rübsamen, M., Roßner, S., Leergaard, T., Kreshuk, A., Puchades, M.A., Bjaalie, J.G., 2019. QUINT: Workflow for Quantification and Spatial Analysis of Features in Histological Images From Rodent Brain. Front. Neuroinform. 13, 75. URL: https://www.frontiersin.org/article/10.3389/fninf.2019.00075/full, doi:10.3389/fninf.2019.00075.

Yonehara, K., Balint, K., Noda, M., Nagel, G., Bamberg, E., Roska, B., 2011. Spatially asymmetric reorganization of inhibition establishes a motion-sensitive circuit. Nature 469, 407–410. URL: https://www.nature.com/articles/nature09711, doi:10.1038/nature09711.

Zhu, Y., Xu, J., Hauswirth, W.W., DeVries, S.H., 2014. Genetically Targeted Binary Labeling of Retinal Neurons. Journal of Neuroscience 34, 7845– 7861. URL: https://www.jneurosci.org/lookup/doi/10.1523/JNEUROSCI.2960-13.2014, doi:10.1523/JNEUROSCI.2960-13.2014.

