## Supplementary Information for "A Knock-In *Igfn1^iCre^* transgenic mouse line provides partial developmental access to type-7 bipolar cells"

Functional neuronal circuits in the vertebrate retina emerge through coordinated developmental events, yet the timeline by which bipolar cells acquire visual feature selectivity remains unclear. A major barrier is the limited genetic access to bipolar subtypes during early postnatal stages. Recent comprehensive transcriptomic study points to Igfn1 as a molecular marker for type-7 bipolar cells (BC 7), a subtype that exhibits direction-selective glutamate releases in adult. Here, we generated an Igfn1iCre knock-in mouse line and characterized Igfn1–positive cell morphology from postnatal day 4(P4) to adult using Cre-dependent tdTomato reporter mice. We found Igfn1–positive cells in the inner retina by P12–P15, predominantly labelling bipolar cells and some amacrine populations. At P15, about 71% of labelled bipolar cells stratified their axons in the S4 sublamina of the inner plexiform layer, consistent with BC 7 morphology. In adult retina, the widespread Igfn1–labelling appears slightly dominated in amacrine cells. To validate these observations, we analysed Igfn1 expression in the Mouse Retina Cell Atlas and confirmed strong Igfn1 enrichment in BC 7 and expression in additional retinal cell types, mirroring experimental results. Overall, these results reveal Igfn1iCre as a potential developmental tool for BC 7 access in the retina.

### S1. Supplementary Figures

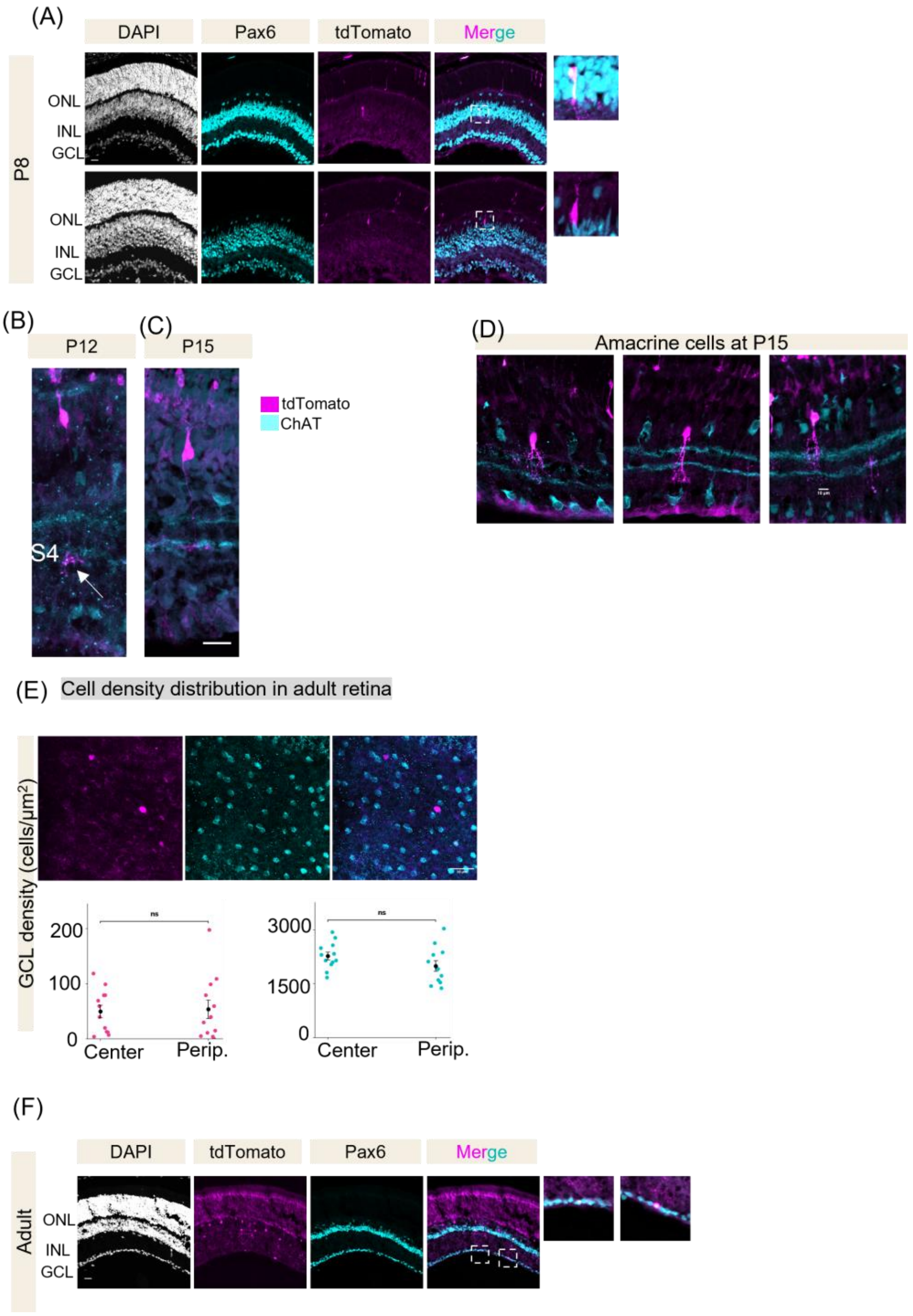


**Figure S1:** (A) Confocal images of P8 retinal sections from *Igfn1* $^{iCre+/-}$; Rosa26-LSL-tdTomato$^{+/-}$ mice stained with pan-AC marker, Pax6. Scale bar, 20 μm. (B–C) Confocal images of representative bipolar cell morphologies at P12, P15. Axon stratification at S4 marks BC 7 identity.Scale bar, 20 μm. (D) Confocal images of representative amacrine cell morphologies at P15. Scale bar, 10 μm. (E) Wholemount image of GCL showing ChAT, tdTomato, and merged channels. Graph shows spatial distribution of tdTomato-positive and ChAT-positive cell densities in the GCL at central and peripheral regions. Student's t-test ($p > 0.05$). (F) Pax6 immunostaining in adult retinal sections. Right side shows high-magnification images and white arrows indicate colocalization. Scale bar, 20 μm.

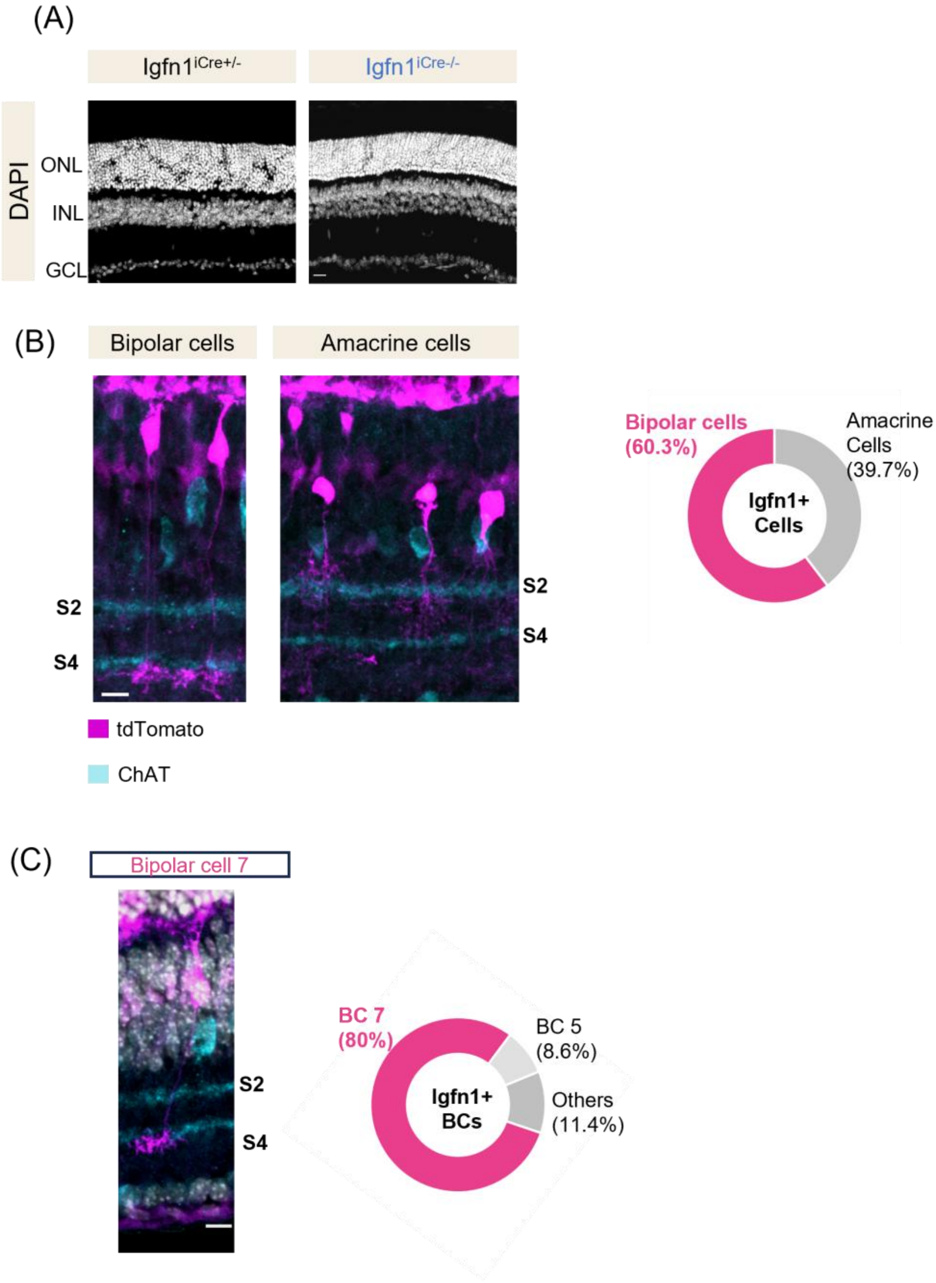


**Figure S2:** (A) DAPI staining of retinal sections from *Igfn1* $^{iCre-/-}$; Rosa26-LSL-tdTomato$^{+/-}$ mice at P15 showing intact lamination of the ONL, INL, and GCL. Scale bar, 20 $\mu$m. (B) Distribution of tdTomato labeled cells in the INL. Approximately 60% of labeled cells were classified as bipolar cells and the remainder as amacrine cells. Scale bar, 10 $\mu$m. (C) Axon terminal stratification of tdTomato-labeled bipolar cells in the inner plexiform layer. Approximately 80% of labeled bipolar cells projected to sublamina S4, consistent with BC 7 identity (n= 58, 3 retinas). Scale bar, 10 $\mu$m.

Type 7 bipolar cell labelled by *Igfn1*$^{iCre}$ transgenic mouse line

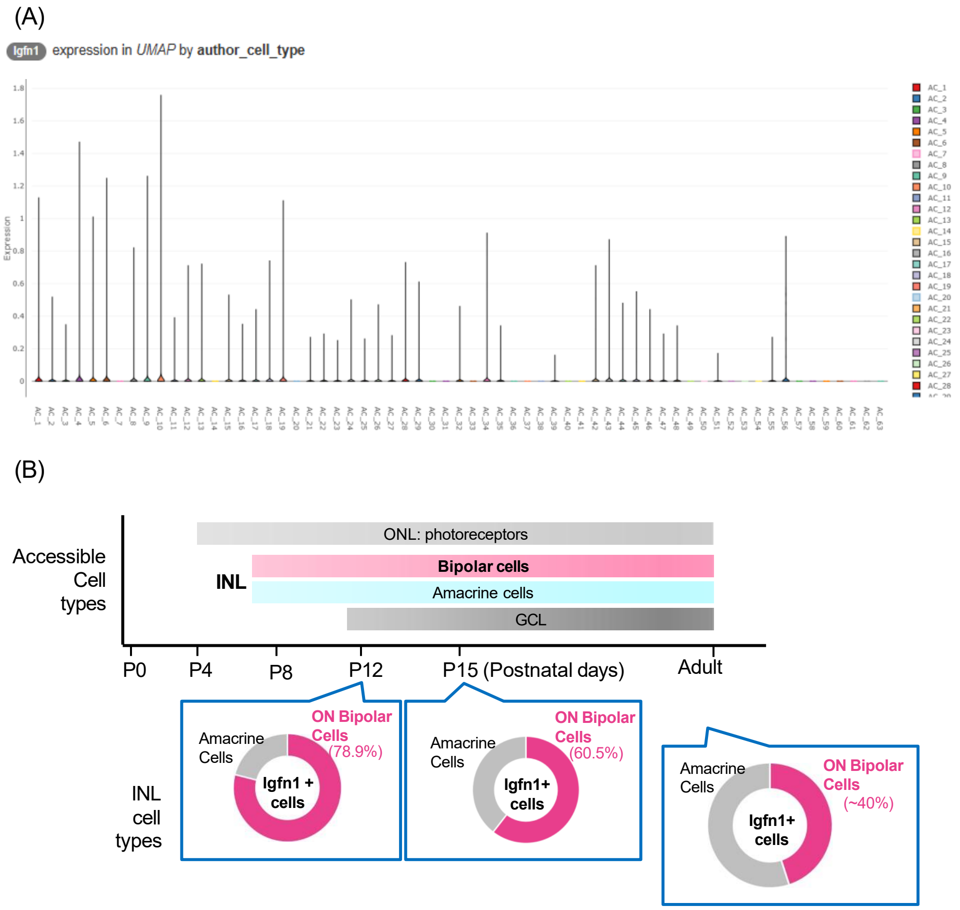


**Figure S3:** (A) *Igfn1* expression across all amacrine cell clusters (B) Summary showing access to retinal cell types by *Igfn1*$^{iCre}$ transgenic line

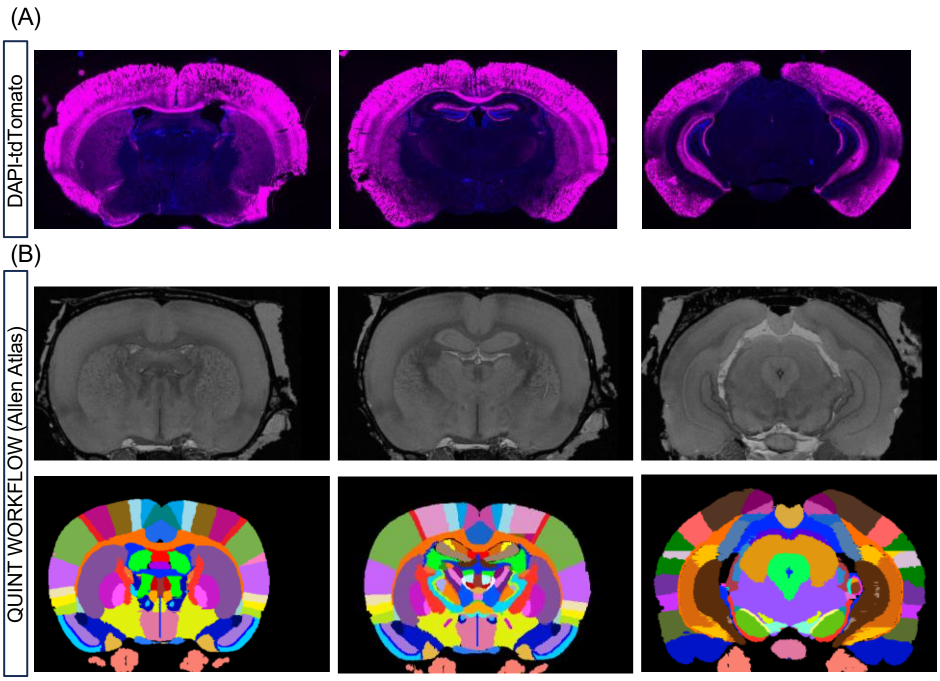


**Figure S4:** (A) DAPI-tdTomato stained merged representative images of *Igfn1* $^{iCre+/-}$; Rosa26-LSL-tdTomato$^{+/-}$ adult mice (B) QUINT workflow softwares- QuickNII and VisuAlign registered to Allen Mouse Brain Atlas were used to find an approximately matching brain section and to determine the annotated regions of interest.
